# Loss of function of *Atrx* leads to activation of alternative lengthening of telomeres in a primary mouse model of sarcoma

**DOI:** 10.1101/2023.11.06.565874

**Authors:** Matthew Pierpoint, Warren Floyd, Amy J. Wisdom, Lixia Luo, Yan Ma, Matthew S. Waitkus, David G. Kirsch

## Abstract

The development of a telomere maintenance mechanism is essential for immortalization in human cancer. While most cancers elongate their telomeres by expression of telomerase, 10-15% of human cancers use a pathway known as alternative lengthening of telomeres (ALT). In this work, we developed a genetically engineered primary mouse model of sarcoma in CAST/EiJ mice which displays multiple molecular features of ALT activation after CRISPR/Cas9 introduction of oncogenic *Kras^G12D^* and loss of function mutations of *Trp53* and *Atrx.* In this model, we demonstrate that the loss of *Atrx* contributes to the development of ALT in an autochthonous tumor, and this process occurs independently of telomerase function by variation of mTR alleles. Furthermore, we find that telomere shortening from the loss of telomerase leads to higher chromosomal instability while loss of *Atrx* and activation of ALT lead to an increase in telomeric instability, telomere sister chromatid exchange, c-circle production, and formation of ALT-associated promyelocytic leukemia bodies (APBs). The development of this primary mouse model of ALT could enable future investigations into therapeutic vulnerabilities of ALT activation and its mechanism of action.

## Introduction

Telomeres are hexanucleotide repetitive elements that are highly conserved to provide stability to the ends of linear chromosomes^1^. During the process of cell division, telomeres are incompletely replicated leading to their progressive shortening, and this is known as the end-replication problem^2^. In order to achieve replicative immortality, most cancers maintain telomere length by activating a mechanism of telomere maintenance, thus preventing telomere erosion and telomere crisis. It is estimated that 85% of cancers maintain telomeres via the activation of canonical telomerase activity. However, approximately 10-15% of cancers use alternative mechanisms to maintain their telomeres^3,4^. These telomerase independent mechanisms of telomere maintenance have been named alternative lengthening of telomeres (ALT). Phenotypic markers of ALT in mammalian cells include abnormally long and heterogeneous telomere length, ALT-associated promyelocytic leukemia bodies (APBs), extrachromosomal telomeric c-circles, and increased frequency of telomere sister chromatid exchange^4,5,6,7^.

Thus far, the genomic determinants of ALT have not been fully elucidated, but the activation of ALT has been strongly correlated with the loss of α-thalassemia/mental retardation, X-linked (ATRX) protein function or expression in multiple subtypes of sarcoma including osteosarcoma, leiomyosarcoma, dedifferentiated liposarcoma, pleomorphic liposarcoma, angiosarcoma, myxofibrosarcoma, and undifferentiated pleomorphic sarcoma^8,9,10,11,12,13^. For patients with these subtypes of sarcoma, the alternative lengthening of telomeres pathway and the loss of ATRX are correlated with shorter progression free survival and worse overall survival of patients^14^*. In vitro* studies have shown that the loss of ATRX in human cells can lead to the development of ALT^15,16,17,18^. Additionally, the reintroduction of ATRX expression represses ALT activity, reinforcing its role as a regulator of telomere dynamics^19,20^.

While traditional mouse models used to study cancer and telomere biology share many commonalities with their human counterparts, they also have key differences. Murine telomerase function is analogous to that of human telomerase, with a mouse telomerase RNA (mTR) which serves as an RNA template for the mouse telomerase reverse transcriptase gene (mTERT) and allows for reverse transcription of telomeres.

However, in contrast to human biology, mice have significantly longer telomeres and have been shown to express telomerase in normal tissues^21^. Despite the prevalence of ALT activation in human cancer, efforts to recapitulate ALT have been largely unsuccessful in primary mouse models of cancer. Additionally, previous studies have demonstrated that, unlike in human cells, telomere maintenance is not essential for the oncogenic transformation of murine cells, independent of their telomere length^22^. In previous work studying telomere shortening after loss of telomerase, CAST/EiJ mice have been used as a model system because they have significantly shorter telomeres than the commonly used 129/SVJ and C57BL/6 strains of laboratory mice^23^. Studies using CAST/EiJ mice with and without genetic loss of mTR have shown that they are able to recapitulate phenotypes of telomere shortening seen in human diseases when telomerase function is lost and that the shortening of telomeres by loss of mTR leads to increased recombination events^24,25^. In this work, we generate and characterize a genetically engineered mouse model of soft tissue sarcoma which recapitulates the alternative lengthening of telomeres pathway after activation of oncogenic *Kras* and loss of function of both *Trp53* and *Atrx*.

## Results

### A CRISPR/Cas9 Primary Model of Sarcoma in CAST/EiJ Mice

For this study, we generated CAST/EiJ mice with constitutively expressed Cas9 (Rosa26-1loxp-Cas9) either with or without functional mTR alleles^22,26^. To initiate primary tumors in these mice, we designed a two-plasmid system to activate oncogenic *Kras^G12D^* and to induce the loss of function of either *Trp53* (CAST KP model) or *Trp53* and *Atrx* (CAST KPA model). In this model, the intramuscular injection and electroporation of plasmids into the right hind limb of mice leads to the formation of spatially and temporally restricted soft tissue sarcomas after approximately 60 days, and this is similar to previously published models^27,28^ (Figure 1A). The first plasmid activates oncogenic *Kras^G12D^* using homology directed repair and the second plasmid contains single guide RNAs (sgRNAs) which induce insertion/deletion (indel) mutations to either *Trp53* alone or *Trp53* and *Atrx* (Figure 1B). In this work, we predict telomerase function by the mTR genotype of mice (Figure 1C). Notably, mTR^+/-^ is haploinsufficient because the loss of functional telomerase activity occurs in both mTR^+/-^ and mTR^-/-^ mice whereas telomerase is able to maintain telomere lengths in mTR^+/+^ mice^29^. Importantly, ATRX is one of the most frequently mutated genes in ALT positive cancers^30^. As a chromatin remodeling protein, ATRX binds with DAXX to deposit histone 3.3 at repetitive elements of DNA, including telomeres (Figure 1D). Previous work has demonstrated that the loss of ATRX leads to telomere dysfunction^31^.

**Figure 1:**
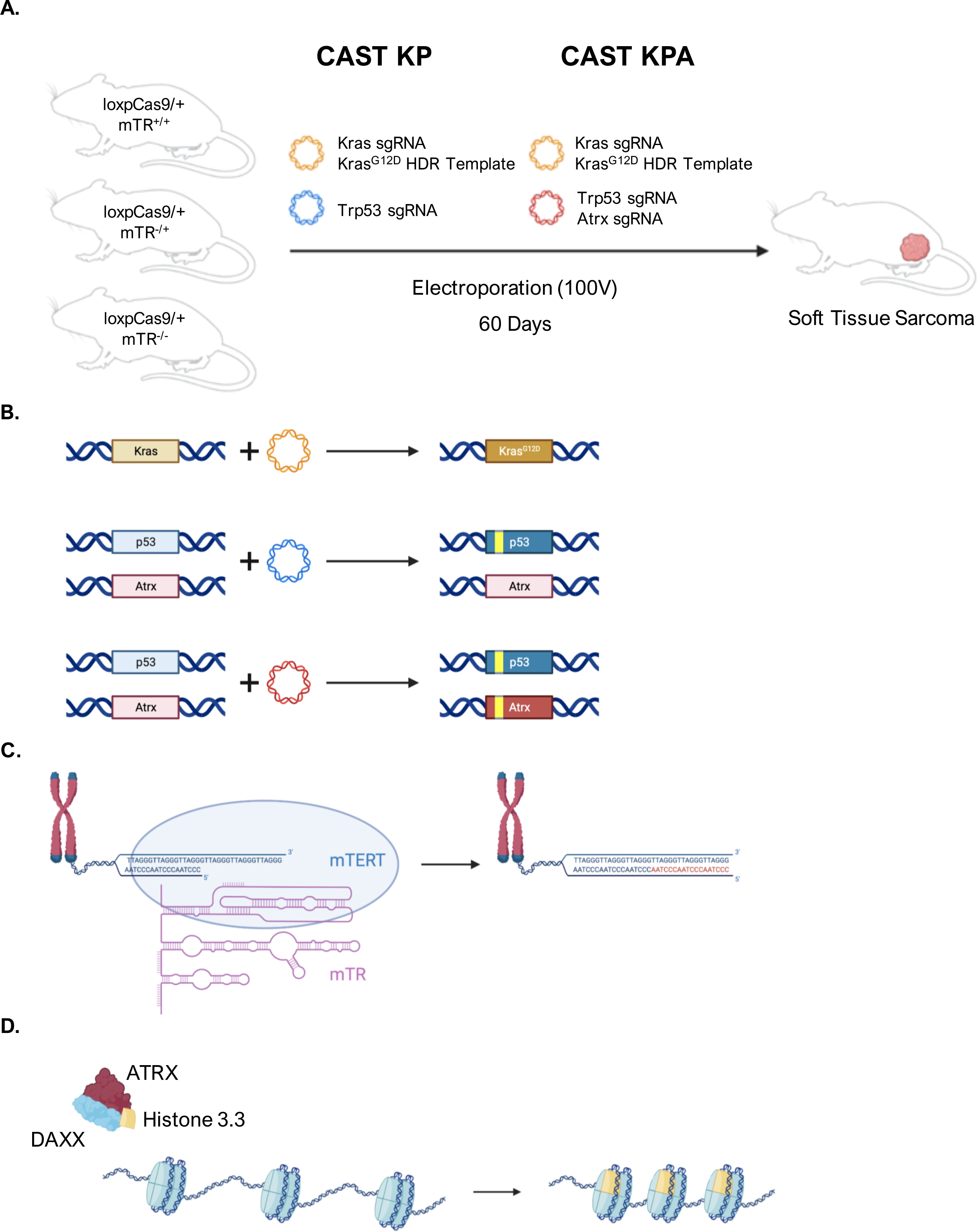
Genetically Engineered Mouse Model of Soft Tissue Sarcoma With Loss of Atrx. (A) CAST/EiJ mice expressing Cas9 (Rosa26-1loxp-Cas9/+) and either wild type mTR (mTR^+/+^), heterozygous mTR loss (mTR^+/-^), or homozygous mTR loss (mTR^-/-^) are injected in the gastrocnemius muscle with a two-plasmid system to induce expression of oncogenic *Kras^G12D^* and biallelic mutation of *Trp53* (CAST KP model), with additional mutation of *Atrx* (CAST KPA model), followed by electroporation of the gastrocnemius muscle. Tumor initiation occurs approximately 60 days after electroporation of the plasmids. (B) Schematic of plasmid activity. The yellow plasmid contains a sgRNA targeting Kras, which is repaired with a homology directed repair template that induces an oncogenic *Kras^G12D^* mutation. The blue plasmid contains a sgRNA targeting *Trp53*, and the red plasmid contains sgRNAs targeting *Trp53* and *Atrx.* (C) mTR is an RNA template for mTERT which is essential for the function of telomerase (D) ATRX as an epigenetic regulator of chromatin. ATRX and DAXX deposit Histone 3.3 at repetitive elements, including telomeres.

### Next-Generation Amplicon Sequencing of Atrx Predicts Expression

To determine the genomic status of *Atrx* after CRISPR/Cas9 editing, we performed targeted next-generation amplicon sequencing of tumor tissue to evaluate the mutation status of each tumor for *Atrx* (Figure 2A). All CAST KP tumors sequenced consisted of entirely wild type *Atrx* sequences. Out of 21 CAST KPA tumors, 14 contained either frameshift or large deletion mutations in *Atrx* (Figure 2B). To test if the results of the amplicon sequencing matched protein expression, we performed immunofluorescence for *Atrx* on three representative primary cell lines. Cell lines with either wild type or non-frameshift sequences demonstrated normal levels of protein expression of *Atrx*, while a cell line with frameshift and structural variant sequences demonstrated loss of *Atrx* protein expression (Figure 2C). Because the loss of *Trp53* and activation of oncogenic *Kras^G12D^* is sufficient to drive tumorigenesis in mice, an additional loss of function mutation of *Atrx* is not required for tumor formation^27^. Therefore, tumors which retain expression of *Atrx* after non-frameshift mutations are classified as CAST KPA* to denote the existence of non-frameshift mutations of *Atrx* and the possible retention of *Atrx* function.

**Figure 2:**
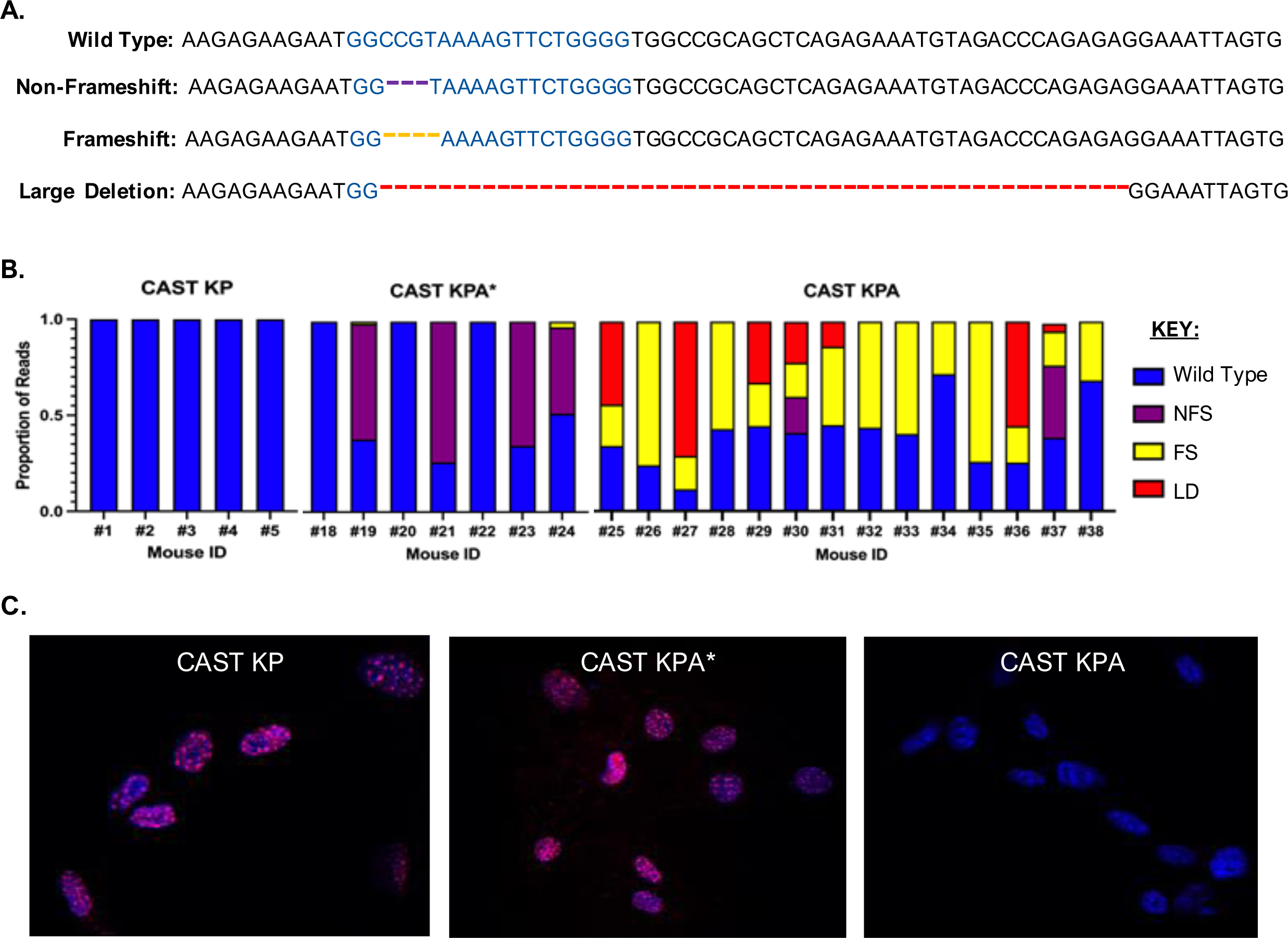
Amplicon Sequencing Predicts *Atrx* Expression. (A) Genomic DNA was extracted from fresh frozen tumors and next generation amplicon sequencing was performed for the site of *Atrx* targeted by CRISPR/Cas9 sgRNA (blue text). Examples of wild-type, non-frameshift (NFS), frameshift (FS), and large deletion (LD) mutation sequences are shown. (B) Graphical representation of the proportion of amplicons in tumors and organization of tumors into wild type CAST KP which did not receive the sgRNA for *Atrx*, CAST KPA* which retain wild type function of *Atrx* after CRISPR/Cas9 editing, and CAST KPA which have loss of function mutations of *Atrx*. (C) Immunofluorescent staining of Atrx (red) and DAPI (blue) of primary cells from CAST KP, CAST KPA*, and CAST KPA.

### Assaying for Alternative Lengthening of Telomeres in Primary Tumors

C-circles are partially single stranded circular telomeric DNAs which are found exclusively in ALT+ tumors and cell lines^6^. The c-circle assay has been used to quantify ALT activity in cell lines and tumors by using phi-29 rolling circle amplification combined with measurement of the resulting products by dot blot or qPCR techniques^6^. In this study, we measured c-circles within genomic DNA from CAST KP, CAST KPA*, and CAST KPA fresh frozen tumors as well as ALT positive U2OS and telomerase positive 143B human cell line controls (Figure 3A). When comparing the relative chemiluminescent signal of the dot blots, these data indicate that the loss of *Atrx* leads to a marked increase in c-circle formation in tumors (Figure 3B). Additionally, non-frameshift mutations in *Atrx* do not lead to the presence of c-circles. These data suggest that loss of function of *Atrx* in this model leads to utilization of the ALT pathway for telomere maintenance.

**Figure 3:**
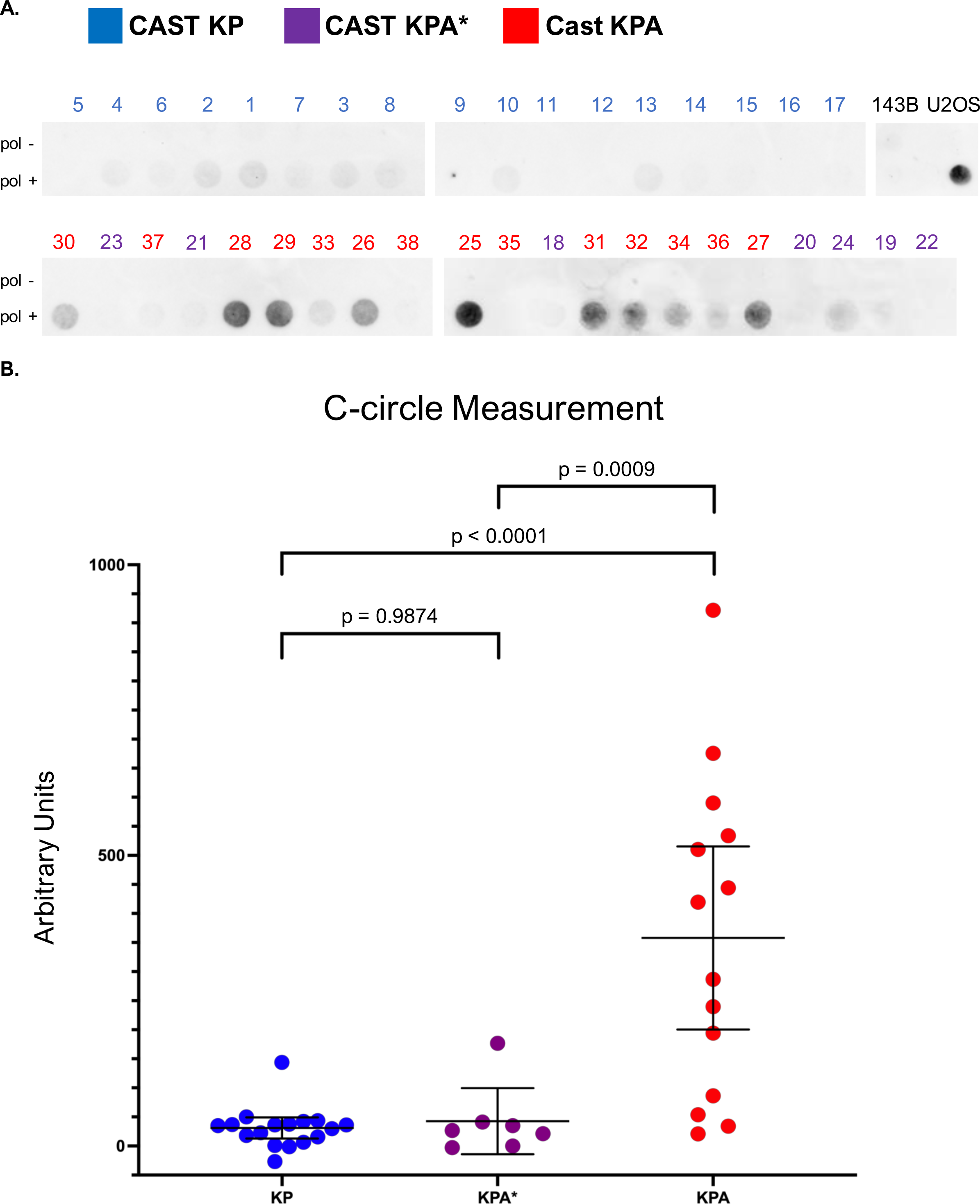
C-circle Measurement in Primary Tumors. (A) Rolling circle amplification of c-circles was performed using DNA extracted from fresh frozen tumors. Chemiluminescent detection of dot blot products with and without the addition of phi29 DNA polymerase from CAST KP (blue), CAST KPA* (purple), and CAST KPA (red) and human control cell lines (black). (B) Graphical representation of the average c-circle content of tumors using a one-way ANOVA with Tukey’s modification.

In addition to c-circles, ALT positive tumors often have unique nuclear PML bodies which colocalize with telomeres. These ALT-associated PML bodies are thought to serve as organizational bodies where the process of telomere maintenance by ALT can occur^5,32^. We performed immunoFISH on frozen tumor sections to analyze the colocalization of PML protein complexes with ultra-bright telomeres (Figure 4). Following imaging, we used a previously established method of analysis to quantify the proportion of APB positive nuclei in each tumor^33^. The presence of ALT-defining characteristics in these CAST KPA tumors, including c-circles and APBs, suggests that this genetically engineered primary mouse model activates the ALT pathway for telomere maintenance.

**Figure 4:**
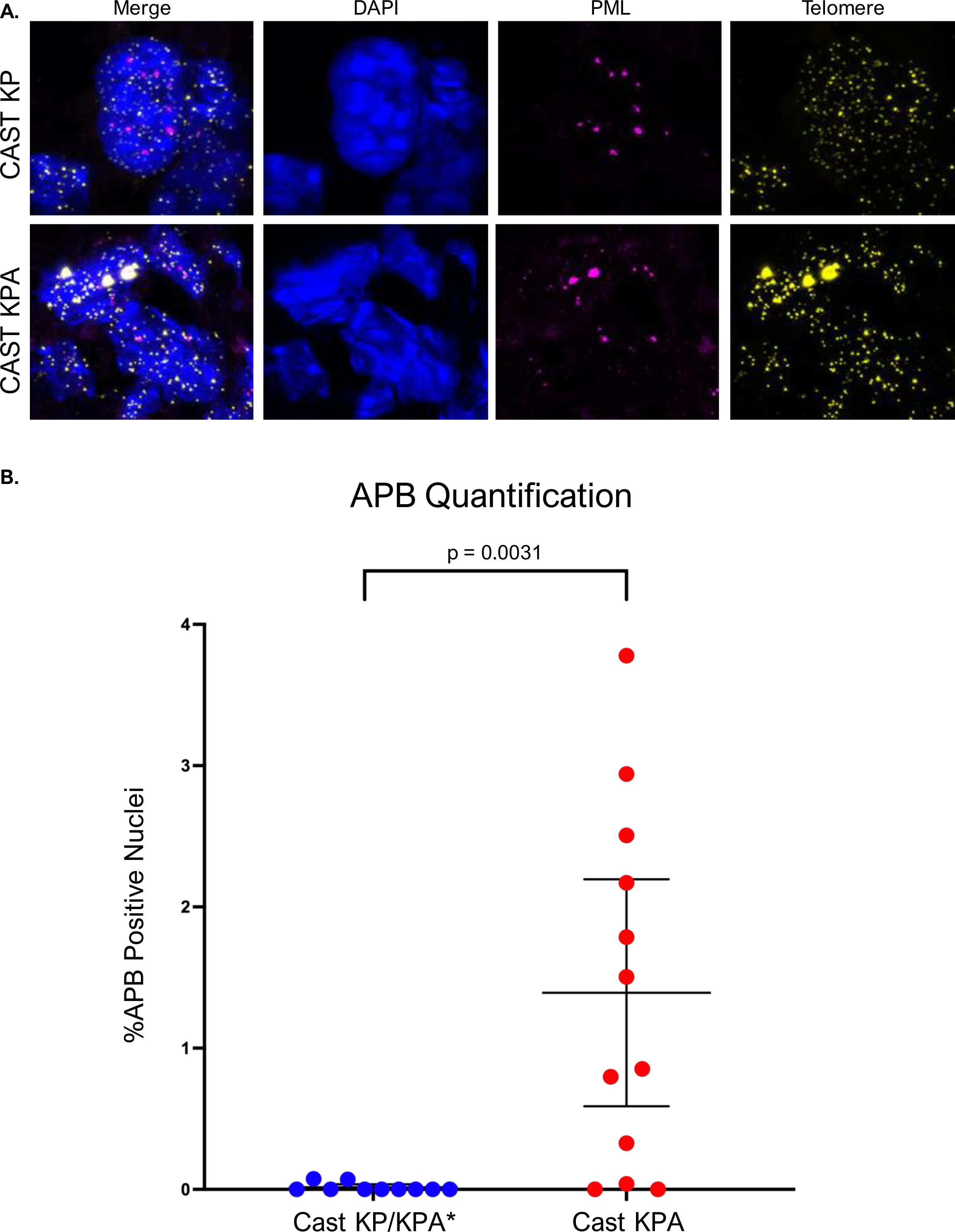
Quantifying ALT-associated PML bodies in Primary Tumors. (A) Representative images of immunoFISH including DAPI (blue), PML (magenta), and telomere (yellow). The colocalization of ultra-bright telomere foci with PML protein is classified as an ALT-associated PML body (APB). (B) Graphical representation of the proportion of APB positive nuclei in individual tumors, compared using a Welch’s t-test.

### Metaphase Analysis of Primary Tumor Cells

To determine the effect of loss of *Atrx* and ALT activation on chromosome structure and telomere phenotypes, we performed metaphase telomere FISH on a panel of primary cell lines derived from CAST KP and CAST KPA tumors with each mTR genotype: mTR^+/+^, mTR^+/-^, and mTR^-/-^ (Figure 5A). We observed that telomere foci decreased in intensity with loss of mTR alleles, and mTR^-/-^ cells had an increase in undetectable telomeres at the end of chromosomes (Figure 5B). Fragile telomeres and chromosome fusions are often a sign of chromosomal instability^34^. In the primary cell lines with loss of function of Atrx, there was an increase in fragile telomeres (Figure 5C). Additionally, in metaphases from cells with loss of both telomerase and *Atrx*, there was an increase in chromosomal fusions compared to cells with loss of telomerase alone (Figure 5D).

**Figure 5:**
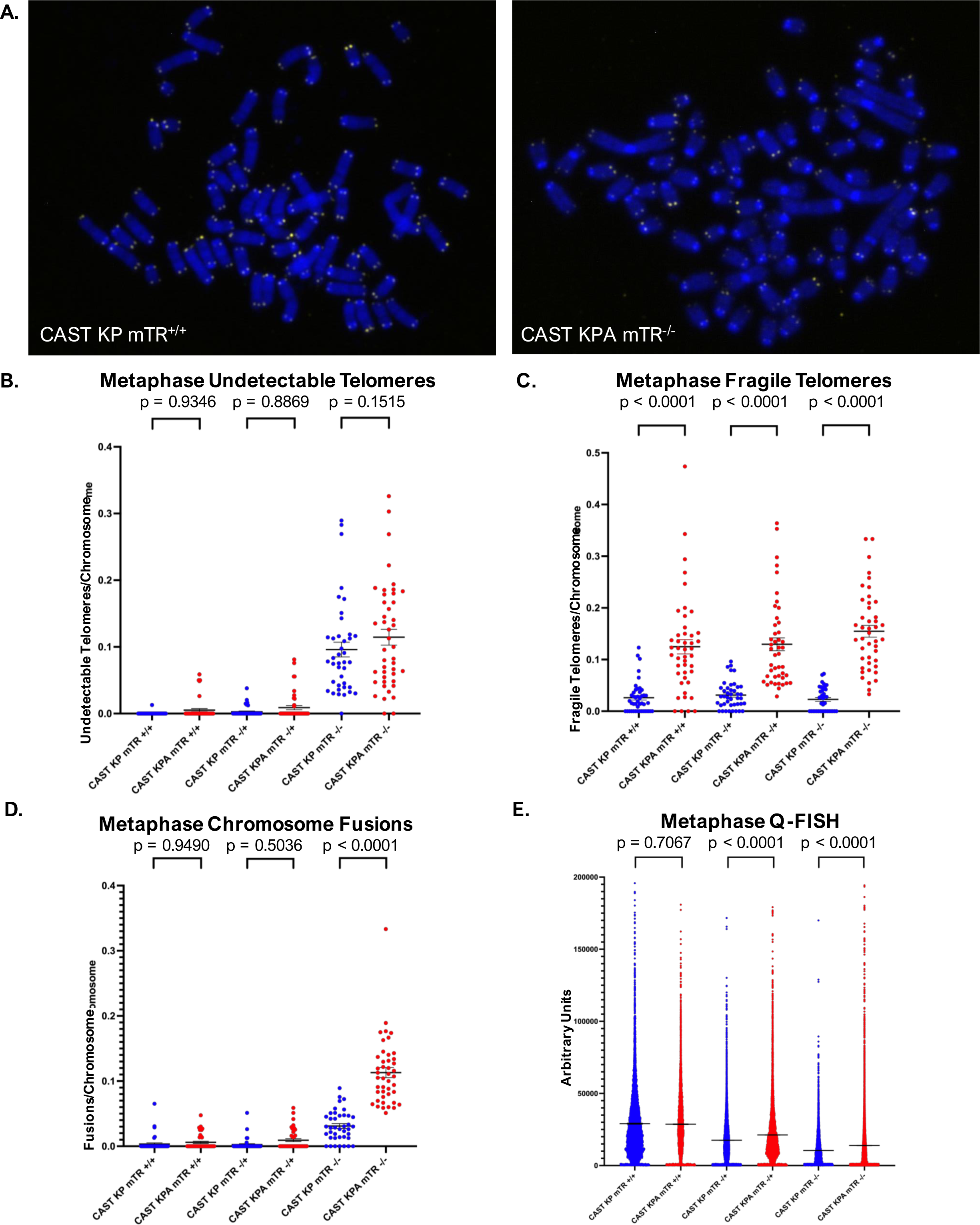
Cytogenetic Analysis of Primary Cell Lines. (A) Metaphase Telomere FISH of primary cell lines, DAPI (blue) and telomere (yellow). (B-D) Graphical representation of (B) undetectable telomeres, (C) fragile telomeres, and (D) chromosome fusions. CAST KP (blue) CAST KPA (red). Statistics performed using one-way ANOVA with Sidak’s multiple comparisons. (E) Q-FISH violin plot of individual telomere fluorescence values from the TFL-Telo program, compared using a one-way ANOVA with Sidak’s multiple comparisons.

To measure the distribution of telomere length in each cell line, we performed quantitative FISH (Q-FISH). To analyze the fluorescent intensity of telomere foci, we used the program TFL-Telo which predicts relative telomere length^35^. In both the mTR^+/-^ and mTR^-/-^ conditions, CAST KPA cell lines had longer average telomere length than CAST KP (Figure 5E). However, in the mTR+/+ condition, we found no significant difference in average telomere length, suggesting that functional telomerase may inhibit the development of telomere heterogeneity in ALT+ cells.

To quantify the number of telomere sister chromatid exchanges (tSCE) in each cell line, we performed chromosome orientation telomere FISH (CO-FISH) (Figure 6A-B). We found a significant increase in tSCE in cell lines derived from tumors with multiple molecular phenotypes of ALT and loss of function of *Atrx* without mTR expression, but we did not observe a difference in tSCE in telomerase positive cells either with or without functional *Atrx* (Figure 6C).

**Figure 6:**
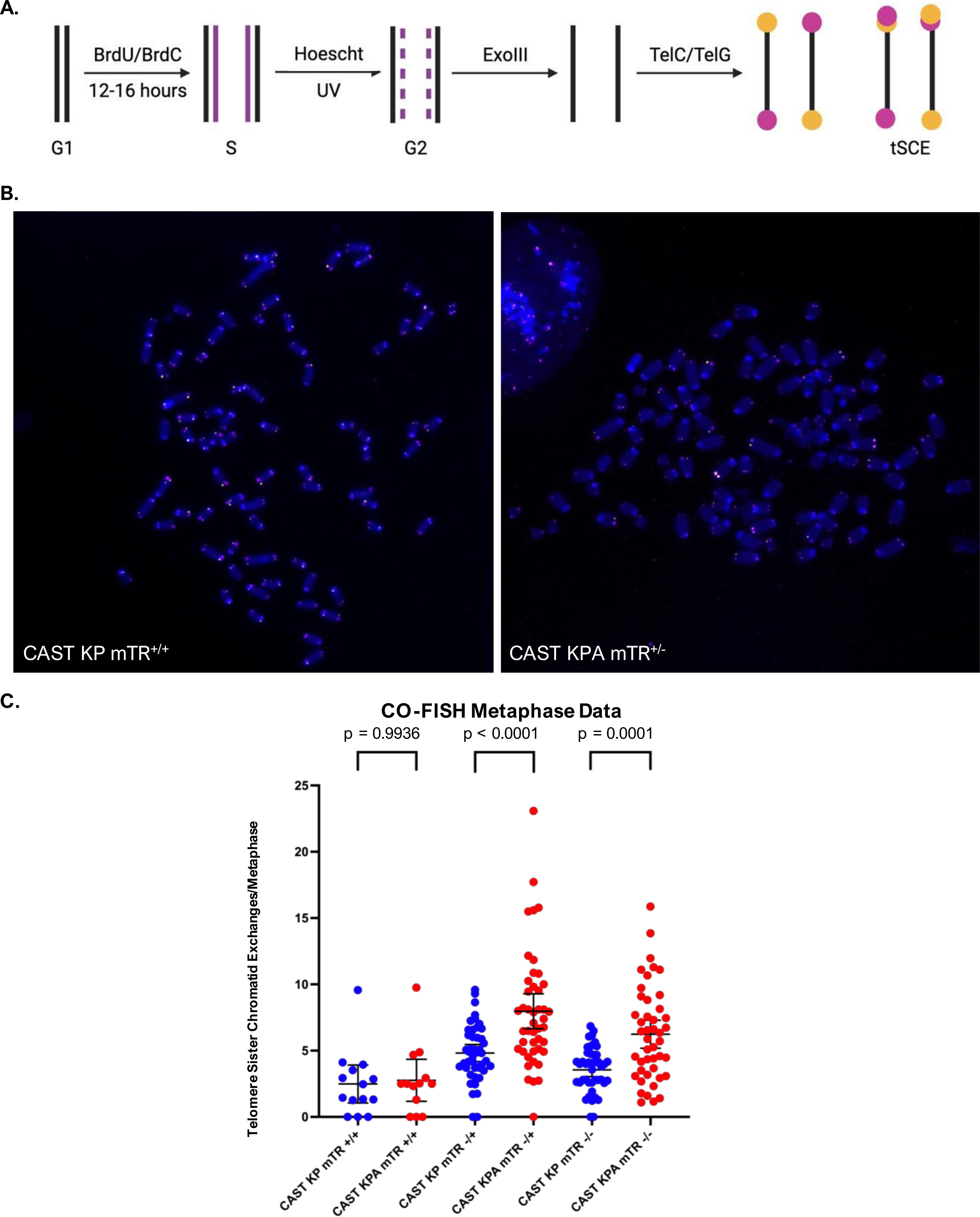
Chromosome Orientation FISH (CO-FISH). (A) Schematic of CO-FISH. (B) Representative images of CO-FISH metaphase performed on primary cell lines. (C) Graphical representation of tSCE quantification in each cell line. Statistics performed using a one-way ANOVA with Sidak’s multiple comparisons.

## Discussion

Despite its prevalence in 10-15% of human cancers, the genomic determinants and specific mechanism of alternative lengthening of telomeres remain unknown. Importantly, the lack of ALT positive autochthonous preclinical models limits the ability to elucidate these mechanisms, to develop targeted therapies for cancers with the alternative lengthening of telomeres pathway, and to investigate the activation of ALT in an immune competent context. Previous *in vitro* work using E14 mouse embryonic stem cells without mTR show that these cells grow for many population doublings before entering telomere crisis and inducing the alternative lengthening of telomeres pathway^36^. While these cells have been shown to extend their telomeres using a template known as mTALT, they do not demonstrate production of c-circles or formation of ALT-associated PML bodies which are characteristic of human ALT positive cancer cells^37^.

In this work, we describe a spatially and temporally restricted primary mouse model of sarcoma which displays multiple molecular phenotypes of ALT. We use a genetically engineered CAST/EiJ mouse strain with shorter telomeres than traditional laboratory mice, with and without loss of telomerase function, to more accurately recapitulate human telomere biology. In this model, the generation of tumors using CRISPR/Cas9 to induce oncogenic *Kras^G12D^* and loss of function of both *Trp53* and *Atrx* leads to sarcomagenesis and to eventual activation of the alternative lengthening of telomeres pathway in a subset of tumors. This primary model will enable the study of the genomic and epigenomic determinants of ALT, the mechanisms by which ALT extends telomeres, and the analysis of therapies for ALT positive tumors.

While the loss of ATRX has been correlated with activation of ALT in human cancers, this study demonstrates a causal relationship between the loss of *Atrx* and the activation of ALT. These findings support a model where the loss of function of *Atrx* drives the formation of ALT but alone may not be sufficient for the activation of ALT *in vivo*.

Interestingly, this work demonstrates that the activation of ALT in this model occurs independently of telomerase function in mice with and without expression of mTR, an essential RNA subunit of telomerase. Generating tumors in mice with mTR^+/+^, mTR^+/-^, and mTR^-/-^ allows us to determine the potential impact of telomerase function on the development of ALT. In this study, we examined the effect of loss of telomerase and loss of *Atrx* function together and separately. We find that loss of telomerase alone leads to telomere shortening and chromosome fusions, while loss of *Atrx* and activation of ALT coincides with an increase in fragile telomeres. Additionally, the combination of loss of telomerase and loss of *Atrx* leads to an increase in tSCEs, chromosomal fusions, and mean telomere length. This telomerase independent recapitulation of ALT phenotypes suggests that the loss of ATRX works in concert with as yet undefined genomic or epigenomic factors to activate ALT in this model. Future work will be needed to investigate these other determinants of ALT and may enable the identification of novel targets for future therapies.

While previous studies have hypothesized that telomerase expression may repress the activation and function of ALT, recent evidence has demonstrated the co-occurence of multiple telomere maintenance mechanisms in human cancer^17,18,38^. The CAST/EiJ KP and KPA mouse models provide additional evidence of alternative lengthening of telomeres in tumors with functional telomerase. Although the telomere restriction fragment assay has been used to show telomere length heterogeneity and lengthening of telomeres considered to be hallmarks of ALT, we find that in telomerase positive cell lines, ALT activation did not impact average telomere length or telomere length distribution, suggesting that the presence of telomerase in these tumors is able to regulate telomere length dynamics independent of ALT activation. However, the function of telomerase does not affect the production of c-circles or the formation of APBs in these tumors. These results indicate that the measurement of telomere length distribution alone is not sufficient to define ALT and that the presence of c-circles and APBs is a more accurate predictor of ALT status in tumors harboring activation of telomerase in the clinic.

One of the limitations of this study is that it does not fully define the mechanism and co-determinants which enable ALT activation in *Atrx*-deficient tumors. While we predict that the loss of epigenetic regulation of telomeres by *Atrx* leads to increased telomere instability and DNA damage that has been theorized to promote ALT, future studies will be needed to precisely delineate the mechanisms and timing by which the loss of *Atrx* leads to activation of the ALT pathway. Because the progressively shorter average telomere length occurs in cell lines with partial (+/-) and full loss (-/-) of mTR, with and without ALT, one possibility for the timeline of ALT activation in this model is that the pathway is activated after significant telomere shortening from cell division with multiple rounds of DNA replication. In this scenario, telomere shortening induces a selection pressure which favors cells activating ALT in a heterogeneous tumor environment. A second possibility is that the activation of ALT occurs as an early event during tumor formation, resulting from the loss of *Atrx* function. While the majority of tumors with loss of function of *Atrx* activated the ALT pathway, a subset of tumors without functional *Atrx* were ALT-, suggesting that the loss of *Atrx* alone is not sufficient to activate ALT in a primary tumor.

While telomerase has been challenging to target therapeutically due to its importance in normal stem cell function leading to toxicity, proteins essential to ALT may be better targets for future treatments as the pathway is exclusive to cancer cells. In mouse xenograft tumors, it has also been found that anti-telomerase therapy has led to cancer regression, followed by tumor recurrence with the development of ALT as a key mechanism of resistance^39^. The prevalence of ALT in cancer and the escape of telomerase positive cancers after telomerase inhibition by gaining ALT function make targeting the ALT pathway an increasingly important area for further research and a promising target for future therapies for many cancer types.

In summary, using this primary mouse model of sarcoma with alternative lengthening of telomeres, we demonstrate that the mutation of *Atrx* facilitates the development of primary cancers with alternative lengthening of telomeres, independent of telomerase function. This work represents the development of a robust ALT-positive primary mouse model of cancer which recapitulates phenotypes observed in human cancer. Development of this model lays the foundation for future research that may allow for improved understanding of the mechanisms of ALT positive cancer and testing ALT-specific therapeutic targets.

## Methods

### Mouse Strains

Mice of mixed background (129SVJ/C57/Bl6) with Rosa26-1loxp-Cas9 and with mTR^-/-^ (B6.Cg-*Terc^tm1Rdp^*/J; Jackson Labs 004132) were crossed with CAST/EiJ mice from Jackson labs (Strain 000928). The mice generated from these crosses were then bred on a pure CAST/EiJ background for five generations. After five generations, mice were electroporated in small batches with littermate controls as they were available. Genotyping was performed by Transnetyx using mouse tails to assess Rosa26-1loxp-Cas9 and mTR genotypes.

### Mouse Tumor Formation

Mice were injected with either KP or KPA plasmids reconstituted at 1ug/ml in half normal saline into the gastrocnemius of the right hind limb using electroporation set at 100V. Mice were checked for tumors after 60 days. The sequences for sgRNA are as follows: *Kras* sgRNA: TACGATAGAGGTAACGCTGC, *Trp53* sgRNA: GTGTAATAGCTCCTGCATGG, *Atrx* sgRNA: CCCCAGAACTTTTACGGCC. The sequence for *Kras^G12D^* HDR is: TGAGAAGTGGACTTTCTTTCTCTGTGGTGAGCTCTCATGAGAAGTGGACTTTCTTGCACCTATGGTTCCCTAACACCCAGTTTAAAGCCTTGGAACTAAAGGACATCACATATAACTGAAATAACTTTCAATATAATTTTCTAATAAATATAAAAATGATATCTTTTTCAAAGCGGCTGGCTGCCGTCCTTTACAAGCGCACGCAGACTGTAGAGCAGCGTTACCTCGATGGTTGGATCATACTCATCCACAAAGTGATTCTGAATTAGCTGTATCGTCAAAGCGCTCTTGCCCACGCCGTCGGCGCCCACGACCACAAGTTTATACTCAGTCATTTTCAGCAGGCCTTACAATAAAAATAATAAAATACAACTAAATTAGAACATGTCTCACACAAGATTATCAAAAACTTTATCAATATTTTAACACTCACCTTTGTGTGTAAAACTCTAAGATATTCCGAATTCAGTGACTACAGATGTACAGAGTAACCTGTAAGTCACTTACAACTTTCTCATCAACACTTAATAAAAATACATGGAAACCAGTACTTTCATCTTCTATCATTTAGCTTGTTGATCTAAACAGCCAGATTACTGTTTTGTAGCAGCTAATGGCTCTCAAAGGAATGTATCATGACTTCACTCAGTACAAATATTCTGCATAGTACGCTATACCCTGTGGACACACCCGCATGAGCTTGTCGACAGCTATCCCAACACCTACCCTTGCGGTATTCTTTGTTGACTGGATATTAAAAGTTAGAAGTTAGGTAGCCTAAGAACATCTGTGTTGGAGCCA

### Cell Line Generation from Primary Tumors

After reaching growth endpoint, tumors were collected from sacrificed mice and digested in dissociation buffer in PBS containing Collagenase Type IV, dispase, and trypsin. Cells were washed in PBS and filtered using a 40um sieve before culture. After five passages, cells were frozen for use in experiments.

### Amplicon Sequencing

Fresh frozen tumors embedded in OCT from CAST KP and CAST KPA mice were cut into 50um slices and stored in Eppendorf Tubes at –20 °C. To prepare tumor tissue for DNA extraction, 1mL of cold PBS was added to each sample on ice. Then, tubes were spun down and residual PBS with dissolved OCT was removed. DNA was extracted using Qiagen DNeasy Blood and Tissue kit. 20ng of DNA was aliquoted from each tumor sample and amplified in a Bio-Rad thermocycler for 35 cycles of PCR using AccuPrime Taq Polymerase with the following primers for *Atrx* exon 8a Forward Primer: GAG ACG GCA ACA GTG GGA CT and Reverse Primer: ACA TTG CAG GGT TGC TTT CTG (IDT). PCR products were sequenced by Genewiz using Amplicon-EZ, and sequences were categorized as wild type (no indel), non-frameshift (indel with size divisible by 3 and less than 50), frameshift (indel with size not divisible by 3 and less than 50), or structural variant (indel with size greater than 50) by alignment using the NCBI BLAST tool.

### Immunofluorescence for Atrx

Cells were seeded on Nunc Lab-Tek II Chambered Coverglass (Thermofischer Scientific, 155379). After 24-48 hours, cells were fixed in 4% paraformaldehyde at room temperature for 10 minutes. After fixation, cells were washed and blocked using the MAXpack Immunostaining Media Kit (Active Motif). Cells were incubated overnight at 4 °C in primary antibody (1:100 Abcam Rabbit anti-Atrx ab97508). After primary incubation, cells were washed 3 x 5 minutes in PBST (PBS + 0.1% Tween 20) and incubated in secondary antibody at room temperature for 1 hour (1:1000 Goat anti-Rabbit IgG Alexa Fluor 488 –cross absorbed). Cells were then washed 3 x 5 minutes in PBST and mounted using Prolong Diamond antifade with DAPI. Representative images were taken of cell lines using the Leica SP5 confocal microscope with consistent settings to allow for comparison.

### C-circle Assay

Fresh frozen tumors embedded in OCT were cut into 20um slices and stored in Eppendorf Tubes at –20 °C. To prepare tumor tissue for DNA extraction, 1mL of cold PBS was added to each sample on ice. Then, tubes were spun down and residual PBS with dissolved OCT was removed. DNA was extracted using QCP DNA extraction described previously^6^. Briefly, while samples and Qiagen protease are placed in ice, QCP lysis buffer is warmed to 56 °C. Qiagen protease is added to QCP lysis buffer at a ratio of 1:20, and 50ul of the resulting lysis buffer is added to each tissue sample. Samples were then incubated at 56 °C for one hour with periodic vortexing and incubated at 70 °C for 20 minutes to inactivate the protease. DNA was quantified using the Qubit HS assay. 1ul containing 40ng of DNA was aliquoted into 9ul of 10mM Tris HCL pH 7.5 in preparation for rolling circle amplification. To perform amplification of c-circles using Phi29 polymerase, 10ul of Master Mix with or without phi29 polymerase was added to each sample and gently mixed. Samples were incubated in a Bio-Rad thermocycler at 30 °C for 8 hours, then heat inactivated at 70 °C for 20 minutes and stored at 11 °C until being collected. 60ul of 2x saline-sodium citrate (SSC) was added to each sample after amplification was complete. To prepare for dot blotting of c-circle products, a nylon membrane was cut to fit the dot blotting apparatus (GE Whatman Minifold I). The nylon membrane was hydrated in water for 5 minutes, then in 2x SSC for 5 minutes. The dot blot apparatus was prepared as described in the manual and bubbles were removed from the nylon membrane. To ensure the vacuum was working properly, 100ul of 2x SSC was loaded into each well and the vacuum was turned on. Then, samples were loaded into each well using a multichannel pipette, with 2x SSC added to empty wells, and the vacuum was turned on. Lastly, 100ul of 2x SSC was added to each well and the vacuum was turned on. While the membrane was still moist, it was crosslinked twice at 454 nm for 1200J using the auto crosslink function (Stratagene UV Stratlinker 1800) before preparing for hybridization. Hybridization of the DIG-telomere probe and chemiluminescent detection was performed according to the protocol from the TeloTAGGG Telomere Length Assay (Roche). Briefly, the nylon membrane was prehybridized in prewarmed DIG Easy Hyb Granules for 60 minutes at 42 °C. Next, the membrane was hybridized with the telomere probe for three hours at 42 °C. The membrane was then washed and blocked before incubation with the Anti-DIG-AP. After washing and incubation in detection buffer, substrate solution was added to activate chemiluminescence. Images were taken using the Chemidoc MP imaging system and the Image Lab software. The c-circle assay was performed on each tumor sample three times using unique adjacent frozen slices, and the results of these experiments were averaged. For each experiment, U2OS was included as a positive control, and 143B was included as a negative control.

### ImmunoFISH

This protocol was adapted from previous methods^32^. Briefly, fresh frozen tumors embedded in OCT were cut into 5um slices and mounted to charged slides before being frozen at –80 °C. Slides were fixed in ice cold methanol for 20 minutes at –20 °C. Slides were then rehydrated using an ethanol series each for 3 minutes at room temperature (100%, 95%, 70%, di H2O). Slides were then washed in 1% Tween 20 for one minute before antigen retrieval using citrate buffer pH 6 by boiling in a rice cooker for 30 minutes. Slides were then cooled for 10 minutes and washed in di H2O. Slides were then dehydrated in (70%, 95%, 100%) ethanol and allowed to air dry. To each slide, 50ul of PNA probe was added, and samples were denatured at 83 °C for 5 minutes. Following this step, slides were protected from light. Slides were incubated in a hybridization chamber for 2 hours at room temperature and subsequently washed in PNA wash buffer 2 x 15 minutes. Slides were washed in PBST 3 x 5 minutes. Then, slides were incubated in primary antibody for PML (1:100 Mouse Anti-PML Antibody, clone 36.1-104) for 45 minutes at room temperature. Next, the slides were washed 3 x 5 minutes in PBST and incubated in secondary antibody (1:100 Goat anti-Mouse IgG Alexa Fluor 647 Cross-absorbed secondary antibody) for 30 minutes at room temperature. Slides were washed 3 x 5 minutes in PBST and once in di H2O before being mounted in Prolong Glass Antifade with DAPI. Slides were stored at room temperature for 24 hours in the dark and one week at 4 °C before imaging using the Leica STED confocal microscope.

During imaging, 10 tiles from each tumor slide were randomly chosen using DAPI staining as a reference for z-stack imaging. ALT-associated PML bodies were quantified in max intensity projection image by manually counting the colocalization of ultra-bright telomere foci and PML bodies. Image names were randomized for blinded analysis. For each quantified APB, it was confirmed that colocalization occurred on a single z-plane image. Representative images for Figure 4 were taken using the 100x/1.4 HCX PL APO OIL DIC WD 90 um objective with 5x Zoom at 4096×4096 resolution and deconvolved using Huygens Essential. All immunoFISH images are displayed as maximum intensity projections using ImageJ.

### Metaphase Telomere FISH

Exponentially growing primary mouse sarcoma cells derived from CAST KP and CAST KPA tumors were incubated in 100ng/mL Colcemid for 1 hour. Cells were then collected using trypsin and incubated in hypotonic solution at 37 °C for 8 minutes. Fixative solution was added dropwise, and cells were fixed overnight at 4 °C. Cells were then resuspended in fresh fixative solution and metaphase spreads were made on ethanol washed slides. Slides were incubated overnight at room temperature. Slides were washed in PBS before fixation in 4% paraformaldehyde at room temperature for 5 minutes. Slides were then washed in PBS 3 x 5 minutes. Metaphase spreads were treated in 250 ug/mL RNase A for 15 minutes at 37 °C in a humidified chamber. Then slides were fixed a second time in 4% paraformaldehyde for 5 minutes at room temperature and washed 3x 5 minutes in PBS. Slides were dehydrated using an ethanol series (70%, 95%, 100%) and allowed to air dry. Chromosomes were incubated in Tel C Cy 3 PNA probe (PNA Bio) at 83 °C for 5 minutes and allowed to hybridize at room temperature in a dark hybridization chamber for 2 hours at room temperature. Slides were washed 2 x 10 minutes in PNA wash A, then 3 x 5 minutes in PNA wash B before a quick wash in di H2O and mounting in Prolong Glass Antifade with DAPI. All fluorescent widefield images were taken using identical settings to allow for comparison.

### Q-FISH

The TFL-Telo program was used to generate brightness estimation of telomeric foci.

### CO-FISH

Primary mouse sarcoma cells derived from CAST KP and CAST KPA tumors were incubated in 10uM 3:1 BrdU/BrdC in growth media for 12-16 hours (7.5uM BrdU, 2.5uM BrdC). Cells were then incubated in fresh media with 100ng/mL Colcemid for 2 hours. Cells were then collected using trypsin and incubated in hypotonic solution at 37 °C for 8 minutes. Fixative solution was added dropwise, and cells were fixed overnight at 4 °C. Cells were then resuspended in fresh fixative solution and metaphase spreads were made on ethanol washed slides. Slides were incubated overnight at room temperature. Chromosomes were then washed in PBS and fixed in 2% paraformaldehyde for 10 minutes at room temperature before washing 3 x 5 minutes in PBS. Slides were then washed in 2x SSC and incubated in 2x SSC with 0.5ug /mL Hoescht stain for 15 minutes in the dark. Slides were then exposed to 365nm UV light for 30 minutes using a Stratalinker and washed in di H2O. DNA labeled with BrdU and BrdC was then digested using Exonuclease III at 37 °C for 30 minutes. Slides were washed in PBS. Next, DNA was denatured by heating at 70 °C for 10 minutes in (70% formamide, 30% 2x SSC). After denaturing, slides were dehydrated using an ethanol series (70%, 95%, 100%) and allowed to air dry. Chromosomes were hybridized to Tel G 488 PNA probe (PNA Bio) and washed in PNA Wash Buffer A before hybridization with Tec C Cy3 PNA probe (PNA Bio) and subsequent washes in PNA Wash A and PNA Wash B. Slides were washed in di H2O and mounted using Prolong Glass Antifade with DAPI. Images were taken using consistent settings on a widefield fluorescent microscope. Telomere sister chromatid exchanges were quantified using signal from only the Tel C probe as it was most consistent between cell lines with different telomere lengths, but representative images of both probes are displayed in the figures after consistent gating of background signal from the Tel G probe using ImageJ.

## Author Contributions

Conceptualization: WF and DGK Methodology: MP, WF, AW, and DGK Formal Analysis: MP, WF Investigation: MP, WF, LL, and YM Wrote the original draft: MP Reviewed and edited the manuscript: MP, WF, AW, MW, and DGK Visualization: MP Supervision: DGK

## Acknowledgements

We thank Dr. Jianguo Huang for providing Rosa26-1loxp-Cas9/+ mice and the Jackson laboratory for providing mTR^-/-^ mice. We thank Dr. Titia de Lange for advice interpreting metaphase telomere FISH phenotypes. We thank Dr. Andrea Daniel for advice in experimental design. Graphics for schematics were created using Biorender.com. This work was supported by 7R35CA197616 from the NCI to DGK.

